# Demand elasticity predicts addiction endophenotypes and the therapeutic efficacy of an orexin/hypocretin-1 receptor antagonist in rats

**DOI:** 10.1101/356709

**Authors:** Morgan H James, Hannah E Bowrey, Colin M Stopper, Gary Aston-Jones

**Author notes:** Corresponding Author: Morgan H. James, Ph.D., 683 Hoes Lane West, Piscataway NJ 08854. E.

## Abstract

Behavioral economics is a powerful, translational approach for measuring drug demand in both humans and animals. Here, we asked if demand for cocaine in rats with limited drug experience could be used to identify individuals most at risk of expressing an addiction phenotype following either long (LgA) or intermittent (IntA) access self-administration schedules, both of which model the transition to uncontrolled drug seeking. Moreover, because the orexin-1 receptor antagonist SB-334867 (SB) is particularly effective at reducing drug-seeking in highly motivated individuals, we asked whether demand measured after prolonged drug experience could predict SB efficacy. Demand elasticity (α) measured immediately following acquisition of cocaine self-administration (‘baseline α’) was positively correlated with α assessed after 2w of LgA or IntA. Baseline α also predicted the magnitude of compulsive responding for cocaine, drug seeking in initial abstinence, and cued reinstatement following LgA, IntA or standard short access (ShA). When demand was measured after LgA, IntA or ShA, α predicted the same addiction endophenotypes predicted by baseline α, as well as primed reinstatement and the emergence of negative emotional mood behavior following abstinence. Post-LgA/IntA/ShA α also predicted the efficacy of SB, such that high demand rats showed greater reductions in motivation for cocaine following SB (10 and 30mg/kg) compared to low demand rats. Together, these findings indicate that α might serve as a behavioral biomarker to predict individuals most likely to progress from controlled to uncontrolled drug use, and to identify individuals most likely to benefit from orexin-based therapies for the treatment of addiction.

## Introduction

A major goal of addiction medicine is to identify those individuals at greatest risk of progressing from controlled to compulsive drug use, such that preventative and therapeutic measures can be targeted in an efficient manner (Blanchard et al., 2009; Belin et al., 2011). To this end, preclinical studies have sought to identify behavioral biomarkers that predict addiction vulnerability. Rodents with high locomotor reactivity to a novel environment, or a high propensity to visit a new environment in a free-choice procedure, exhibit a greater tendency to exhibit addiction-like behaviors when given the opportunity to self-administer drugs of abuse (Davis et al., 2008; Blanchard et al., 2009; Belin et al., 2011; Flagel et al., 2016). Although these models provide a useful tool for examining addiction vulnerability in rats, the extent to which these phenotyping procedures can be applied to clinical populations is unclear.

Behavioral economics (BE) has emerged as a promising cross-species paradigm for determining addiction propensity (Bentzley et al., 2013; Bentzley et al., 2014; MacKillop et al., 2018). A fundamental construct of BE is demand, or the amount of commodity that is sought or consumed at a given price. Demand for drugs can be measured via a hypothetical purchasing task in humans (MacKillop et al., 2008; MacKillop et al., 2010; MacKillop et al., 2018) or an operant task in rodents (Oleson et al., 2011; Bentzley et al., 2013; Bentzley et al., 2014). Demand elasticity (denoted as alpha; α) refers to the extent to which demand is sensitive to changes in price, and can be modeled in both humans and rodents using identical mathematical approaches (Hursh & Silberberg, 2008; Bentzley et al., 2013; Bentzley et al., 2014). In humans, α predicts several clinical outcomes across several drugs of abuse, including cocaine (MacKillop et al., 2008; MacKillop et al., 2010; MacKillop et al., 2012; Bruner & Johnson, 2014; MacKillop et al., 2018). In animals, we and others have shown that α is correlated with a range of addiction-like behaviors, including compulsive drug taking, drug-seeking during initial abstinence, and reinstatement propensity (Bentzley et al., 2014). However, in many of these studies, α was determined in subjects already meeting the diagnostic criteria for addiction (human studies) or following extensive drug exposure (rat studies). Ideally, an addiction biomarker could be measured early in the addiction cycle (eg. during recreational use) and be used to predict individuals most at risk of progressing to more problematic drug use behavior. Thus, here we sought to determine whether α, as assessed in rats after only limited drug exposure, can predict those individuals most at risk of transitioning to compulsive drug seekers after more drug exposure. To do this, we analyzed data from a number of studies carried out in our laboratory whereby ‘baseline α’ was measured in rats immediately following the acquisition of cocaine self-administration behavior but prior to rats being exposed to either long or intermittent self-administration schedules, both of which model the transition to uncontrolled drug seeking (Ahmed & Koob, 1998; Zimmer et al., 2012; Allain et al., 2015; Kawa et al., 2016). We report that baseline α predicts those individuals that exhibit greater economic demand, compulsive seeking, and reinstatement liability following long or intermittent access to produce uncontrolled drug seeking. Thus, α may serve as a useful screening tool to identify patients that would most benefit from early intervention strategies.

Although a general biomarker might allow for preventative strategies to be more effective and efficient, there will undoubtedly remain a population of individuals who make the transition to uncontrolled drug use and therefore require intervention. To this end, a biomarker that predicts how an individual might respond to a specific pharmacological treatment would be very useful. A large number of studies now point to the hypothalamic orexin (hypocretin) system as a promising target for pharmacotherapies designed to reduce craving and drug-seeking behavior (James et al., 2017; Walker & Lawrence, 2017). Interestingly, compounds that block signaling at the orexin-1 receptor (OXR1) are particularly efficacious at blocking drug taking in individuals with high motivation for drug (Lawrence et al., 2006; Moorman & Aston-Jones, 2009; Jupp et al., 2011; Bentzley & Aston-Jones, 2015; Lopez et al., 2016; Moorman et al., 2017). Thus, in a second set of analyses, we analyzed data from several studies carried out in our laboratory where α for cocaine was determined prior to treatment with the OXR1 antagonist SB-334867. We show that the anti-addiction effects of SB are stronger in animals with higher economic demand for cocaine, indicating that α could also be used as a biomarker to direct pharmacological treatment in a clinical setting.

## Materials and Methods

### Subjects

Rats (Sprague Dawley, male ~300g, 6-8 weeks of age upon arrival; Charles River, Raleigh, NC and Kingston, NY) were single-housed under a reverse 12:12h light cycle in a temperature- and humidity-controlled facility. Food and water were provided ad libitum. For all behavioral experiments, rats were tested during their dark (active) cycle at the same time each day. All experiments were conducted between January 2015 and January 2018 and the data contributed to several manuscripts in submission (James et al., in submission) or preparation. All experiments were conducted in accordance with procedures established by the Institutional Animal Care and Use Committee of Rutgers University, and were conducted according to specifications of the Guide for the care and use of laboratory animals (National Research Council, 2011).

### Drugs

Cocaine HCl powder was provided by the National Institute of Drug Abuse (NIDA; Research Triangle Park, NC, USA) and was dissolved in sterile saline. SB-334867 (SB) [1-(2-methylbenzoxazol-6-yl)-3-[1,5]naphthyridin-4-yl urea hydrochloride; provided by NIDA] was suspended in 2% dimethylsulfoxide and 10% 2-hydroxypropyl-b-cyclodextrin (Sigma) in sterile water; 0, 10 or 30 mg/kg was given in a volume of 4 ml/kg (i.p.) 30 min prior to testing. SB has 50-fold selectivity for OX1R over OX2R and 100-fold selectivity over approximately 50 other molecular targets (Porter et al., 2001; Smart et al., 2001).

### Jugular catheter surgery and cocaine self-administration training

Jugular catheter surgery was performed as previously described (James et al., 2016; McGlinchey et al., 2016; James et al., 2018). Briefly, in anesthetized animals, chronic indwelling catheters were inserted into the right jugular vein and exited the body via a port between the scapulae. Rats were allowed 1w recovery before commencing self-administration training.

Rats were trained to self-administer cocaine in sound-attenuated operant boxes with two retractable levers (Med Associates) under a fixed-ratio 1 (FR1) schedule. The house light was illuminated and active lever presses resulted in a delivery of 3.6s infusion of cocaine (dose; i.v.) paired with a tone and light stimulus above the active lever. A 20s time out period was signaled following each infusion by the turning off of the house light, during which time active lever responses no longer resulted in an infusion of cocaine. Responses on the inactive lever had no scheduled consequences but were recorded. Rats trained in 2h sessions for at least 6d, until they achieved stable cocaine consumption (>20 infusions, <20% variability) across 3d.

### Economic demand procedure

We used a within-session behavioral economics paradigm, previously described elsewhere (Bentzley et al., 2013; Bentzley et al., 2014). Briefly, rats received access to decreasing doses of cocaine in successive 10 min intervals on a quarter logarithmic scale (383.5, 215.6, 121.3, 68.2, 38.3, 21.6, 12.1, 6.8, 3.8, 2.2 and 1.2 μg/infusion), over 110 min sessions. For the duration of each infusion, the houselight was turned off and light+tone cues were presented; active lever responses did not result in further infusions during this period. Demand curves were fit to each animal’s data from each session using our lab’s focused-fitting approach (Bentzley et al., 2013) where the values α and Q_0_ in the exponential demand equation (Hursh & Silberberg, 2008) were manipulated to minimize the residual sum of squares. Curves were fit to data points that occurred up to two points following the maximum price (Pmax) at which the subject maintained its preferred level of drug consumption, as determined by a <25% decrease in brain cocaine concentration, with the omission of the first (‘loading’) data point. α is an index of demand elasticity, an inverse measure of motivation for drug, and Q_0_ estimates consumption at null effort (Hursh & Silberberg, 2008).

### Intermittent, Long and Short Access self-administration paradigms

Animals were randomly allocated to one of three access conditions (ShA, LgA or IntA), in which ShA and LgA rats were given continuous access to cocaine for 1h or 6h, respectively, for 14d (Ahmed & Koob, 1998). The sessions were matched to self-administration FR1 training where the cocaine dose, schedule of reinforcement, light and tone presentation and time out periods during these sessions was identical. IntA rats were repeatedly given 5 min access to cocaine followed by a 25 min time out period (lights out, levers retracted) over a 6h session (total 1h access to cocaine). As per previous studies (Zimmer et al., 2012), cocaine infusions were brief (1s; 0.055mg) and were paired with light+tone cues. No timeout periods were imposed during the 5min epochs when cocaine was available, apart from the time during which the pump was active. The return of cocaine availability following each 25min period of cocaine non-availability was signaled by the presentation of the light+tone cues for 5s and a single priming injection of cocaine (1s, 0.055mg) before the levers were reinserted into the operant chamber. Some animals included in the SB analyses were trained on the ShA and IntA paradigms as above, however training sessions were separated by 1-4d (average of 1 session every 2d).

### Test of compulsive (punished) responding for cocaine

A punished responding task (Bentzley et al., 2014) was used to test rats for compulsive drug-seeking behavior. Animals were allowed to adjust to a relatively high cocaine infusion dose (0.38mg cocaine in a 2.6 sec iv infusion) over at least three sessions, and until the number of infusions earned per session was stable (<20% variation). Throughout the test session, an identical cocaine dose was held constant, but infusions were paired with shocks of increasing amplitude. Footshock was omitted for the first two 10-min bins, however, commencing in the third bin, shocks were delivered during each cocaine infusion and these increased in amplitude every 10min on a tenth-log10 scale: 0.13, 0.16, 0.20, 0.25, 0.32, 0.40, 0.50, 0.63, and 0.79 milliamps (mA). Throughout the session, shock resistance was defined as the maximum cumulative charge in millicoulombs (mC) a rat self-administered in any one bin.

### Extinction and Reinstatement Testing

All extinction training and reinstatement testing occurred in 2h sessions in identical operant chambers to that used for the self-administration training. In extinction training, lever responding was extinguished in daily sessions in which the active lever responses no longer resulted in a cocaine infusion, nor light+tone presentation. Extinction training continued for at least 7d and until rats met a criterion of <25 active lever responses over 3 consecutive days. We tested cued reinstatement behavior where light+tone cues were presented in the absence of cocaine. This occurred once at the beginning of the session and then again in response to active lever responses. To assess primed reinstatement behavior rats were given a priming dose of cocaine (10mg/kg, i.p.) immediately prior to being placed in the operant chamber. Animals were tested under extinction conditions. Between reinstatement tests, rats received at least 2 additional extinction sessions, and continued until they met a criteria of <25 active lever presses over 2d.

### Open Field Test

To evaluate anxiety-like behavior, rats were positioned in a clear acrylic open field apparatus (42cm x 42cm x 30cm). The apparatus was equipped with SuperFlex monitors (Omintech Electronics Inc, Columbus, OH) containing two16 × 16 infrared light beam arrays: one to measure horizontal activity and another to measure vertical activity. Locomotor activity was recorded by Fusion SuperFlex software. The amount of time spent within an 8×8 square matrix in the center of the chamber was determined over a period of 15min.

### Saccharin Preference Test

Rats were habituated to 2 bottles over 2 consecutive days. During this period, 1 bottle containing a sweet 0.1% saccharin solution and the other containing water, were placed in the rat’s home cage. The location of the bottles were counterbalanced and were interchanged every 12h. The following day at the peak of the animals’ active period (noon), the 2 bottles were re-introduced to the cage for a total of 2h, with the locations of each bottle swapped after 1h (test day 1). This same procedure was repeated the following day (test day 2). Preference for saccharin was determined by calculating the amount of saccharin consumed as a percentage of total liquid intake. Saccharin preference scores were averaged across both test days for each animal.

### Forced Swim Test

Animals were placed in a cylindrical Plexiglas tank (38.5 cm high × 30.5 cm diameter; INSTECH) filled with warm water (25°C). Pretesting occurred on the first day for 15 min and testing occurred 24h later for 5 min. A digital video camera mounted over the tank recorded behaviors during both FSTs. Each animal was scored off-line on a computer monitor by an observer blind to the experimental conditions of the animals for swimming, climbing and immobility behaviors, as described elsewhere (Gonzalez & Aston-Jones, 2008).

### Experimental overview

*Experiment 1*: Animals underwent FR1 self-administration training (mean=6.36 days, SD=1.22 days, range 6-11 days) before being tested on the behavioral economics procedure to determine baseline demand for cocaine. Animals were then given LgA, IntA or ShA to cocaine for 14d, after which they were tested on the BE paradigm again to examine the effect of these differential access conditions on cocaine demand. Previous studies demonstrated that baseline demand measures remain stable over weeks (Bentzley et al., 2014), a finding confirmed here in ShA animals (see below). Here, we focus our analyses on demand scores from the first session following ShA/LgA/IntA, as this timepoint is associated with the most pronounced changes in the LgA group (James et al., in submission); using these data allows us to best assess the demand paradigm as a predictor of LgA-induced changes in addiction behavior. Rats underwent testing for compulsive (punished) responding, and then extinction training, followed by tests of cued- or primed-reinstatement behavior. A subgroup of rats was then subjected to 4w homecage abstinence, during which time they were left undisturbed except for weekly cage changes. These rats were then tested on the SPT, OFT and FST (in this order), with at least 5d separating each test. These results are from experiments reported with other analyses in another manuscript (James et al., in submission).

*Experiment 2*: Animals were assessed for baseline demand, subjected to the ShA/LgA/IntA procedures, and then re-tested for demand as above. Following stabilization of demand (~7d; denoted ‘post-LgA/IntA/ShA α’), rats were tested for cocaine demand following injections of SB (0, 10, 30mg; i.p., order counterbalanced). Approximately half of animals (n=44) included in these analyses were also included as part of Experiment 1 – SB testing took place prior to testing for compulsive responding.

### Statistical analyses

All analyses were carried out using Prism GraphPad and SigmaPlot V14.0. For all analyses, α values were normalized using log transformation. All correlations depict transformed α values; in some instances (**Fig 1g-i**), raw α values are presented for clarity. Changes in demand following LgA, IntA or ShA across populations were compared using paired-samples t-tests. Changes in demand in high versus low demand populations were assessed first by a median split of baseline α values (lower α values = high demand group; higher α values = low demand group), and then analyzed using a 2 group (high demand, low demand) x 2 time (pre-, post-LgA/IntA/ShA) mixed model ANOVA with Bonferroni post-hoc comparisons. All correlations were carried out using tests of Pearson’s correlation coefficients. Across all correlations, 3 data points were excluded as they were identified as outliers by Prism auto-detection software. For SB analyses, overall change in α values following SB treatment was assessed across the population using paired-samples t-tests relative to vehicle treatment. The population was then divided into high-, intermediate- and low-demand groups using a quartile split of α values under vehicle; efficacy of SB (calculated as percentage change in α values relative to vehicle) was then compared across both doses using a repeated-measures ANOVA. An α value of 0.05 was adopted for all tests. All analyses used two tailed tests.

**Figure 1.**
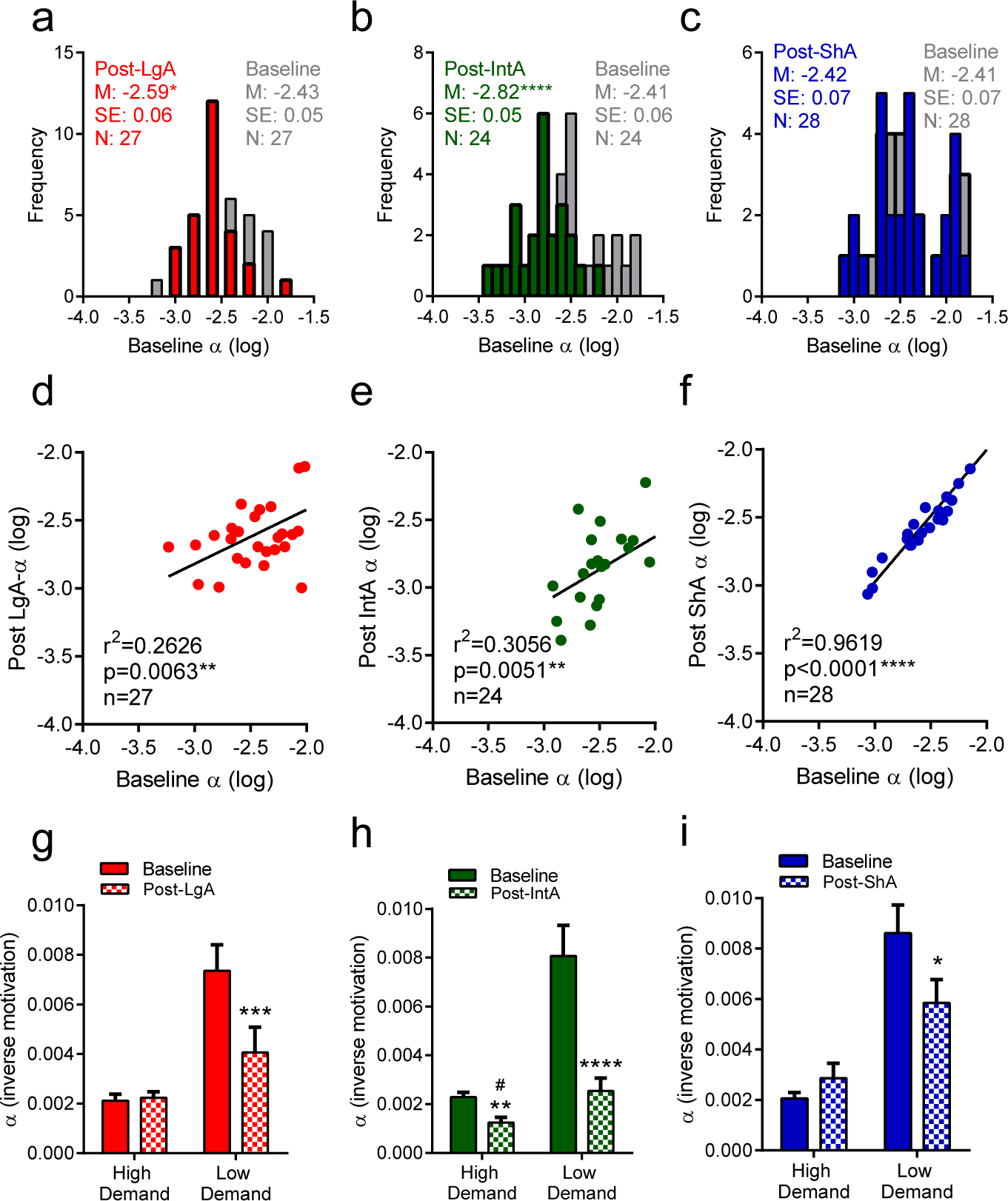
Baseline demand predicts motivation for cocaine following long (LgA) or intermittent (IntA) access. (**a, b**) Frequency histogram of α values following limited cocaine experience (‘baseline’; gray) and following LgA (red; **a**) or IntA (green; **b**) in the same animals. α values are significantly shifted towards the left following LgA or IntA, indicating lower demand elasticity (higher motivation) for cocaine. Note that this shift is more pronounced for InTA than for LgA animals. (**c**) Frequency histogram depicting no change in α values from pre-(gray) to post-(blue) ShA. (**d, e**) Baseline α values predicted α values following LgA (**a**) or IntA (**b**), indicating that animals with the highest demand for cocaine at baseline went on to have the highest demand for cocaine following a transition to compulsive drug seeking. (**f**) Pre- and post-ShA α values were also correlated; because there was no shift in demand following ShA, this relationship indicates that demand is highly stable across time (14d) under normal limited access conditions. (**g**) Animals were split into high-(**HD**) and low-(**LD**) demand groups based on a median split of baseline α values. LgA induced a significant decrease in α values (increased motivation) in LD animals only. Demand tended to be lower in HD animals compared to LD animals following LgA. ***P<0.001 vs. pre-LgA LD. (**h**) IntA was associated with a significant reduction in α values (increased motivation) in both HD and LD animals. Following IntA, α values were lower in HD animals compared to LD animals. # P<0.05 vs. post-IntA LD. **P<0.01 vs. pre-IntA HD. **** P<0.0001 vs. pre-IntA LD animals. (**i**) ShA was associated with a modest decrease in α values in LD animals only. *P<0.05 vs. pre-ShA LD.

## Results

### Baseline economic demand is an effective biomarker for predicting those individuals most likely to exhibit greater addiction-like behavior following LgA or IntA

Following LgA or IntA, rats exhibited a significant decrease in α compared to ‘baseline’ α, demonstrated by a significant leftward shift in the distribution of α values following LgA (t_26_=2.633, P=0.0140; **Figure 1a**) and an even greater shift following IntA (t_23_=6.928, P<0.0001; **Figure 1b**). This reveals an increase in demand (motivation) for cocaine following LgA or IntA (A more thorough comparison of changes in demand following LgA and IntA is described in James et al., in submission). There was no change in demand values following ShA (t_28_=0.6392, P=0.5281; **Figure 1c**). We and others have also reported that these shifts in motivation are associated with the expression of other addiction-relevant behaviors, including increased compulsive (punished) responding for cocaine, increased drug seeking during initial abstinence and enhanced relapse vulnerability. Thus, the LgA and IntA models serve as effective models to study the transition from controlled to compulsive drug seeking behavior.

To examine whether the LgA- or IntA-induced transition to a more motivated state could be predicted by baseline α values, we correlated these two indices. Baseline α values were positively correlated with α values following both LgA (R^2^ =0.2626, P=0.0063; **Figure 1d**) or IntA (R^2^ =0.3056, P=0.0051; **Figure 1e**), indicating that those animals that exhibit greater demand for cocaine following only limited drug exposure are more likely to transition to a higher-demand state following LgA or IntA. Despite there being no change in α values following ShA, we saw a strong correlation between pre- and post-ShA α values (R^2^ =0.9619, P<0.0001; **Figure 1f**), indicating that demand elasticity is extremely stable across time (in this case 2 weeks) in individuals with limited drug experience, pointing to the reliability of the BE approach as a behavioral biomarker.

To further probe the predictive nature of baseline α values, we carried out a median split on baseline α values across all access groups to derive ‘high risk’ (high demand; HD) and ‘low risk’ (low demand; LD) groups, as might be done clinically. Interestingly, HD rats did not exhibit an overall change in demand following LgA, whereas LgA was associated with a significant decrease in α (increase in motivation) in LD rats (group x time interaction F_1,25_=15.97, P=0.0005). Post-LgA α values tended to be lower in HD compared to LD rats (P=0.1070; **Figure 1g**), indicating that those individuals identified as ‘at risk’ at baseline may continue to exhibit a higher motivation for cocaine following extended cocaine access. In IntA animals, both HD (P=0.0016) and LD (P<0.0001) rats exhibited a significant change in α values (main effect of ‘time’ F_1,22_=54.12, P<0.0001). Those rats identified as HD at baseline had lower α (higher motivation) than baseline LD animals following IntA (P=0.0345; **Figure 1h**). Interestingly, there was a small decrease in α following ShA in the LD group (group x time interaction F_1,26_=6.833, P=0.0147, Bonferroni post-hoc comparison P=0.0304), indicating that even limited access to cocaine can increase motivation for cocaine in animals that initially exhibit low demand for drug early in the addiction cycle (**Figure 1i**).

### Baseline economic demand predicts some addiction-like behaviors following LgA or IntA

Next, we tested whether baseline α values also predict other DSM-V-relevant addiction behaviors following significant drug exposure (i.e., 2 weeks of LgA, IntA or ShA). Baseline α values were negatively correlated with the maximum electric charge animals accepted to maintain stable brain-cocaine concentrations (R^2^ =0.2105, P=0.0161; **Figure 2a**), indicating that animals with higher baseline demand for cocaine exhibited greater levels of compulsive drug seeking following LgA, IntA or ShA. Baseline α values were also negatively correlated with the amount of drug seeking (responses on the active lever) on the first day of extinction (R^2^ =0.1339, P=0.0013; **Figure 2b**) or cued reinstatement of extinguished responding (R^2^ =0.2498, P=0.0016; **Figure 2c**), but did not predict primed reinstatement responding (R^2^ =0.0710, P=0.1548; **Figure 2d**). Baseline α values did not predict negative emotional-like behavior on the saccharin preference (R^2^ =0.0715, P=0.2996; **Figure 2e**) or forced swim (R^2^ =0.3269, P=0.5754; **Figure 2f**) tests, which were conducted following 4 weeks of homecage abstinence.

**Figure 2.**
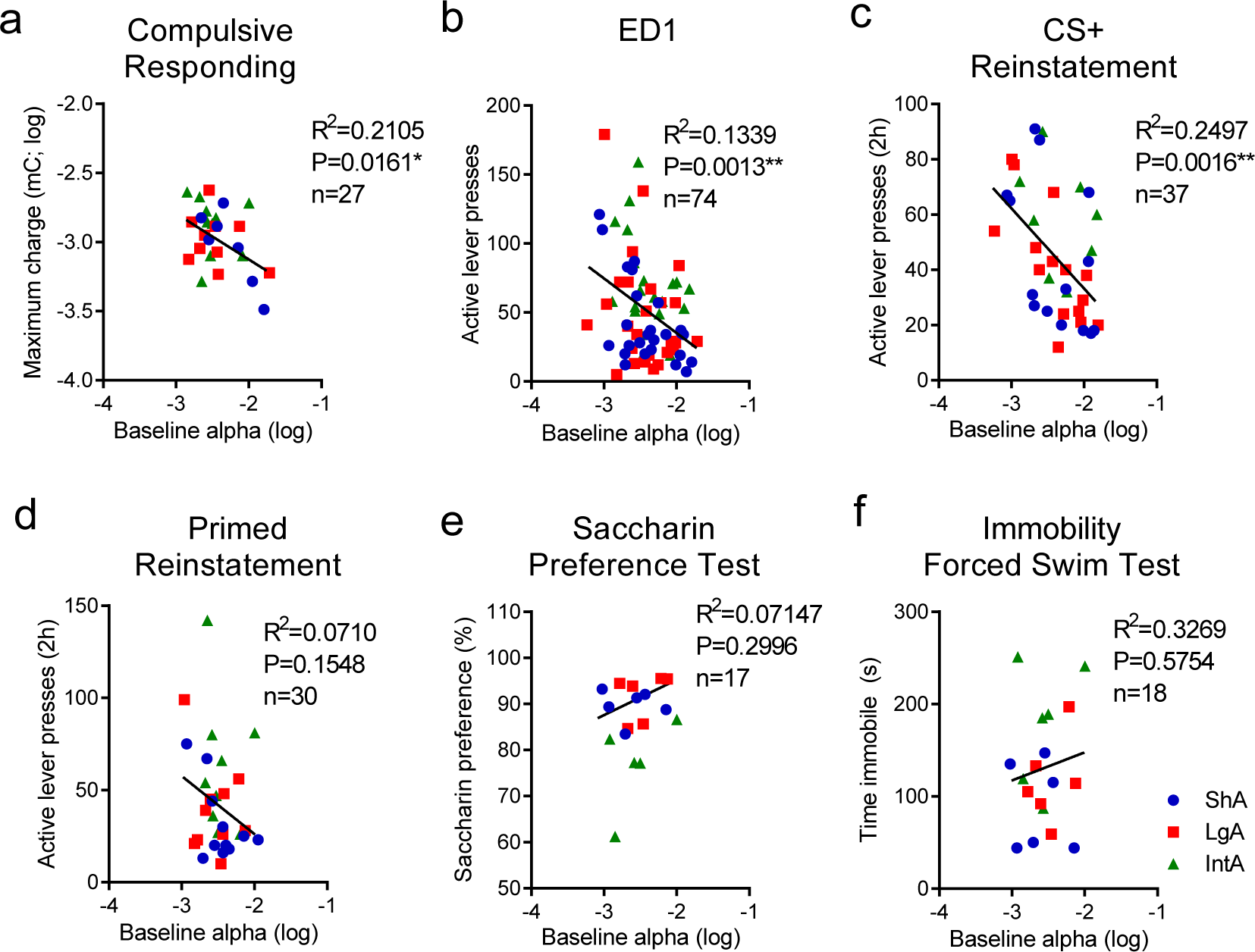
Baseline demand predicts compulsive responding for drug and cued reinstatement behavior following LgA, IntA or ShA. (**a**) Baseline α values were correlated with compulsive responding following LgA (red), IntA (green) or ShA (blue) training, as assessed by the maximum charge animals were willing to accept in millicoulombs (mC) in an FR1 task where cocaine infusions were paired with footshocks of escalating intensity. Low α (higher motivation) was associated with higher compulsive responding. (**b**) Baseline α values were also correlated with drug seeking during initial abstinence (extinction day 1; ED1) following LgA, IntA or ShA. (**c**) Baseline α values also predicted the extent to which animals exhibited reinstatement of extinguished drug seeking elicited by drug associated conditioned stimuli (CS+) following LgA, IntA or ShA. (**d-f**) Baseline α values did not predict cocaine primed reinstatement (**d**), saccharin preference (**e**) or time spent immobile on the forced swim test (**f**) following LgA, IntA or ShA.

### Economic demand following LgA or IntA serves as a strong biomarker for addiction-like behaviors and negative emotional behavior following abstinence

Next, we tested whether α values derived from BE testing after LgA, IntA or ShA provide a better prediction of drug-relevant behaviors than baseline α values. Compared to baseline α values, there was a stronger negative correlation between post-LgA/IntA/ShA α values and compulsive responding (R^2^ =0.3158, p=0.0023; **Figure 3a**) or drug seeking during initial abstinence (R^2^ =0.3822, p<0.0001; **Figure 3b**). Post-LgA/IntA/ShA α values were also highly predictive of both cued (R^2^ =0.2793, p=0.008; **Figure 3c**) and primed (R2=0.2588, p=0.0041; **Figure 3d**) reinstatement behavior. Moreover, unlike baseline α values (described immediately above), post-LgA/IntA/ShA α values were positively correlated with saccharin preference (R^2^ =0.4777, p=0.0021; **Figure 3e**) and negatively correlated with immobility on the forced swim test (R^2^ =0.2845, p=0.0026; **Figure 3f**), indicating that animals with higher motivation for cocaine following LgA/IntA/ShA exhibited greater anhedonia and despair, respectively, following forced abstinence.

**Figure 3.**
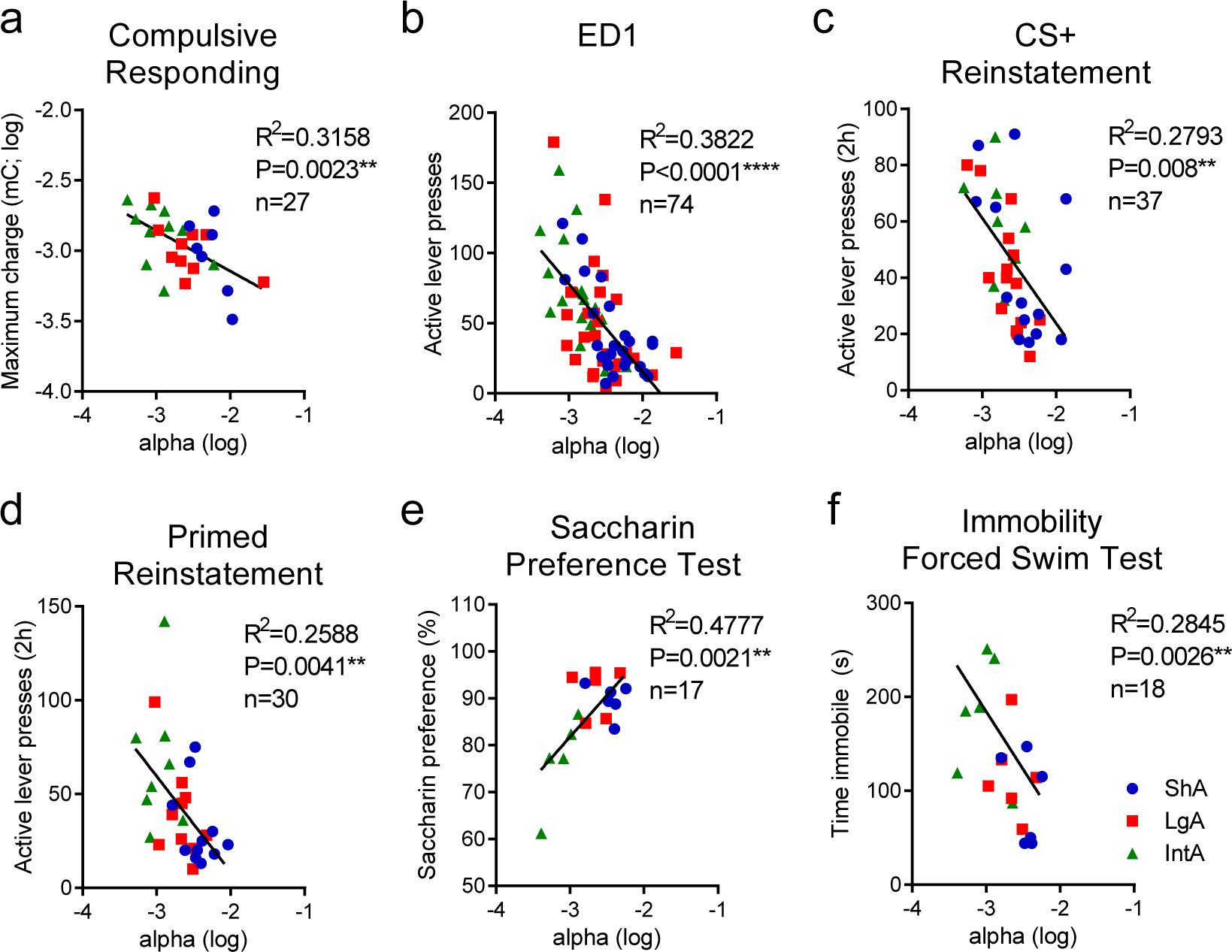
Economic demand for cocaine following LgA, IntA or ShA predicts addiction-like behavior and the emergence of negative emotional behavior following abstinence. (**a-d**) Post-LgA/IntA/ShA α values were negatively correlated with measures of compulsive responding (**a**), drug seeking on the first day of abstinence (**b**), cued reinstatement (**c**) and primed reinstatement, such that animals with lower α values (higher demand) scored higher on each of these measures. (**e**) Post-LgA/IntA/ShA α values were positively correlated with the degree to which animals showed a preference for saccharin on the saccharin preference test following 4 weeks of home cage abstinence. Thus, lower α values (higher demand) were associated with higher withdrawal-induced anhedonia-like behavior. (**f**) Post-LgA/IntA/ShA α values were negative correlated with the amount of time spent immobile in the forced swim test following 4 weeks of home cage abstinence. Thus, lower α values (higher demand) were associated with higher withdrawal-induced depression-like behavior. All correlations reflect Pearson’s correlation coefficients. **P<0.01. ****P<0.0001.

### Economic demand following LgA or IntA predicts the anti-addiction efficacy of the orexin-1 receptor antagonist SB-334867

We previously demonstrated that SB reduces economic demand for cocaine. Other evidence points to SB being particularly effective at reducing cocaine-seeking in high demand animals. Thus, here we tested whether the efficacy of SB at reducing cocaine demand (at two doses; 10 and 30mg/kg) can be predicted by post-LgA/IntA/ShA α values. Across a large population of animals (SB10, n=82; SB30 n=87) animals, both SB10 (t_81_=3.376, P=0.0011; **Figure 4a**) and SB30 (t_86_=5.232, P<0.0001; **Figure 4b**) were associated with a significant rightward shift in α values, indicating greater elasticity in demand for cocaine (lower motivation). Interestingly, there was a strong negative relationship between post-LgA/IntA/ShA α values and the efficacy of either SB10 (R^2^ =0.1508, P=0.003; **Figure 4c**) or SB30 (R^2^ =0.1364, P=0.0004; **Figure 4d**), indicating that animals with greater demand for cocaine (lower α values) exhibited the greatest change in demand elasticity following treatment with both doses of SB.

**Figure 4.**
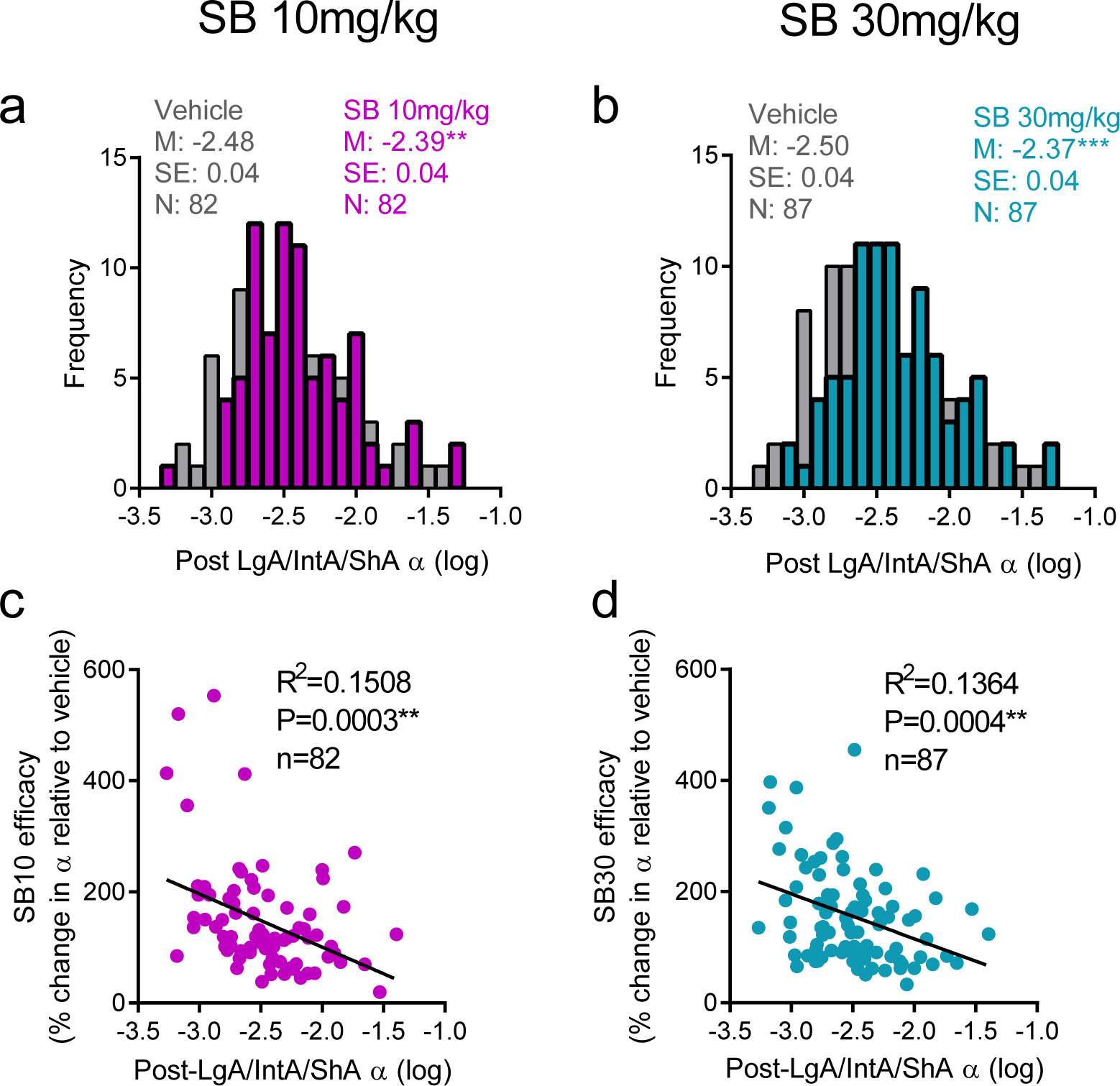
The efficacy of SB in reducing motivation for cocaine is predicted by α values following LgA, IntA or ShA. (**a, b**) Frequency histogram of α values (log transformed) following vehicle (gray), SB10 (purple; a) or SB30 (blue; **b**) treatment. Both SB10 and SB30 treatment were associated with a significant rightward shift in α values compared to vehicle treatment, reflecting a decrease in motivation for cocaine. (**c**) The extent to which SB10 (**c**) or SB30 (**d**) increased α values was negatively correlated with α values following LgA/IntA/ShA, indicating that animals with the highest demand for cocaine (low α values) were more susceptible to the anti-addiction properties of SB.

To further explore this relationship, we divided the total population by their post-LgA/IntA/ShA α values to identify HD (top quartile), intermediate demand (ID; middle two quartiles) and LD (bottom quartile) groups; we then assessed the efficacy of SB (10 or 30 mg/kg, ip) across these different populations. In HD animals, SB10 and SB30 were equally efficacious in reducing demand (increasing α for cocaine (F_2,59_=7.353, P=0.002; **Figure 5a**). In ID animals, both SB10 and SB30 treatment was associated with a significant increase in demand elasticity (decreased motivation; F_2,132_=9.999, P<0.001) and this effect was stronger for SB30 (p<0.001 vs. vehicle; **Figure 5b**). In LD animals, neither SB10 nor SB30 treatment was associated with a change in α (F_2,62_=0.445, P=0.644).

**Figure 5.**
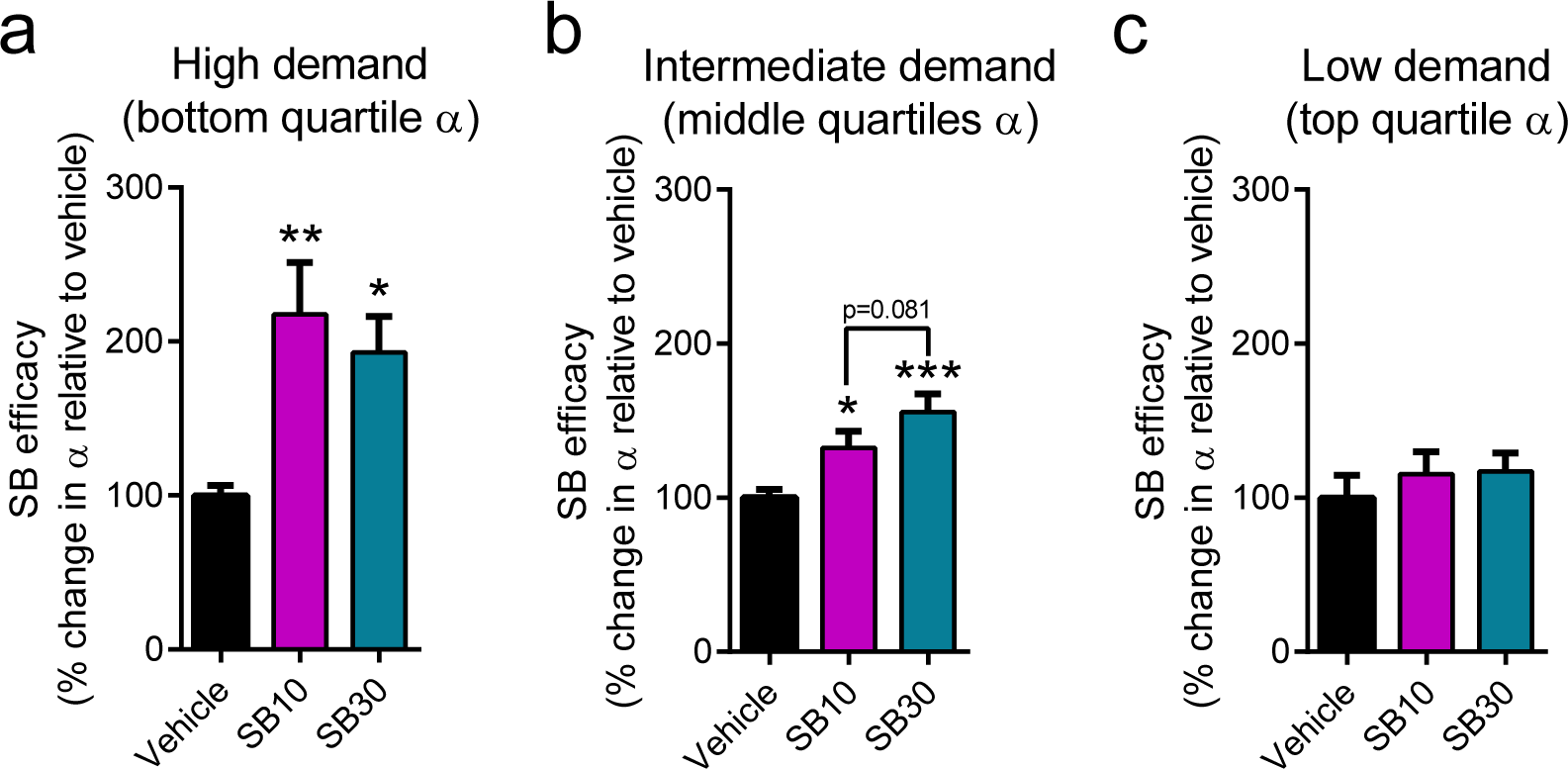
SB differentially affects motivation for cocaine in high-, intermediate- and low-demand animals. Animals were divided into high-, intermediate- and low-demand (HD, ID, LD) groups based on a quartile split of demand values derived from BE testing under vehicle conditions. (**a**) In HD animals, both SB10 (P=0.003) and SB30 (p=0.010) treatments significantly increased α values (decreased motivation) relative to vehicle treatment. There was no significant difference between the efficacy of SB10 and SB30 (P=0.493) in these animals. (**b**) In ID animals, SB10 (P=0.022) and SB30 (P<0.001) also increased α values (decreased motivation). Despite not reaching significance, there was a strong trend towards this effect being stronger for SB30 compared to SB10 (P=0.081) in these animals. (**c**) In LD animals, neither SB10 or SB30 affected α values. Data represents mean ± SEM. All tests represent post-hoc Holm-Sidak comparisons. *p<0.05. **p<0.01. ***p<0.001.

## Discussion

We report three key findings relating to the use of economic demand as a behavioral biomarker for addiction. First, we show that baseline α (demand elasticity) measured after only limited drug exposure predicts the extent to which rats will exhibit excessive motivation for cocaine following cocaine access paradigms (LgA or InTA) that promote the transition to uncontrolled drug use. Moreover, this ‘baseline’ measure of economic demand also predicts an individual’s propensity to exhibit compulsive (punished) drug-seeking, as well as seeking during abstinence or in response to drug stimuli, following LgA or InTA. Second, we show that a reassessment of α following these cocaine abuse paradigms provides an even stronger biomarker for addiction-related endophenotypes, including the emergence of negative emotional behavior following abstinence. Finally, α values strongly predict the extent to which the orexin-1 antagonist, SB, reduces motivation for cocaine, such that SB is more efficacious in high demand (low α) animals but has only limited utility in low demand (high α) animals. Together, these data indicate that economic demand analysis, which is readily applied in humans, may serve as a powerful clinical tool to identify individuals most at risk of progressing from recreational to compulsive drug use, and to identify those most likely to show therapeutic benefit from treatment with an orexin-1 receptor antagonist.

### Demand elasticity as a biomarker for addiction propensity

We previously reported a strong relationship between economic demand and several addiction-related behaviors in rats, including compulsive drug taking and relapse-like behavior (Bentzley et al., 2014). Similarly, measures of demand derived from a hypothetical purchasing task predicts real-world cocaine use in cocaine-dependent patients (Bruner & Johnson, 2014). Parallel findings have been found in studies involving heavy tobacco (MacKillop et al., 2008; Murphy et al., 2011; MacKillop et al., 2012) and alcohol (Murphy & MacKillop, 2006; Murphy et al., 2009; MacKillop et al., 2010; Gray & MacKillop, 2014) users in relation to demand and severity of substance misuse and dependence. Here, we extend these findings to show that economic demand assessment may also be used as a preemptive diagnostic tool to identify individuals most at risk of developing substance use disorder. Rats with a high baseline demand for cocaine (assessed after only limited drug exposure) went on to exhibit greater motivation (lower α values) following LgA or IntA, and there was a strong correlation between baseline demand and the extent to which rats exhibited compulsive drug seeking and cued reinstatement following LgA or IntA. This was despite the greatest relative changes in demand occurring in individuals identified as having low baseline demand for cocaine; indeed only a modest change in demand was observed in HD animals after IntA, whereas no change occurred in HD animals following LgA. Interestingly, LD rats exhibited a modest increase in motivation for cocaine (lower α values) after ShA, indicating that even individuals identified as having low risk of cocaine abuse are susceptible to the motivation-enhancing properties of limited but extended cocaine use.

Consistent with previous reports in animals (Bentzley et al., 2014) and clinical populations (MacKillop et al., 2008; MacKillop et al., 2010; MacKillop et al., 2012), α values derived after the transition to compulsive drug use are an even more robust biomarker of addiction endophenotypes. Indeed, post-LgA/IntA/ShA α values predicted compulsive drug seeking, seeking during initial abstinence and cued reinstatement of extinguished seeking to a stronger extent than baseline α values. In addition, post-LgA/IntA/ShA α values predicted primed reinstatement levels and the expression of anhedonia- and despair-like behavior. This may indicate that although trait factors contribute substantially to addiction-like vulnerability, ultimately this vulnerability reflects combination of both trait and state factors (O’Brien, 2008).

### Comparison with other behavioral biomarkers of addiction

Many studies have identified different patterns of drug-related behavior in animals that exhibit differential reactivity and/or preference to novel environments. For example, rats that exhibit greater locomotor reactivity to a novel inescapable environment (high responder phenotype; HR) learn to self-administer cocaine at a faster rate, have higher rates of drug intake, and are more reactive to the locomotor activating effects of psychostimulants, compared to their low responder (LR) littermates (Piazza et al., 1989; Davis et al., 2008; Blanchard et al., 2009). Following extended cocaine access, HR rats exhibit persistent cocaine seeking (seeking during a non-drug-available period) and increased reinstatement of extinguished cocaine seeking; these rats also exhibit distinct epigenetic modifications in the nucleus accumbens core that are not observed in LR rats (Flagel et al., 2016). An alternative model identifies rats that develop a strong conditioned place preference for a novel environment (high-novelty-preferring phenotype; HNP) (Belin et al., 2011). These HNP rats show a greater severity of cocaine addiction-like behavior across a number of DSM-V-relevant criteria, including persistence in drug seeking (measured by drug seeking during periods when drug is unavailable), compulsive (punished) responding for drug, and motivation for drug (measured on a progressive ratio schedule) (Belin et al., 2011).

Efforts to ‘reverse translate’ the HR/HNP models to clinical populations using tests of unconditioned motoric activity have had some success, however the link between exploratory behavior in humans and psychiatric disorders, including addiction, remains to be fully understood (Young et al., 2016). In contrast, the behavioral economics approach described here provides a direct mathematical link between addiction-like behavior in rats and humans (Hursh & Silberberg, 2008; Bentzley et al., 2013; Bentzley et al., 2014), so that preclinical findings using this approach can be readily translated to the clinic. An interesting feature of HNP/HR models is that they can predict addiction-like behavior prior to drug use. However, individuals most in need of preventative interventions are likely to present to relevant health care providers following some level of recreational drug use, perhaps at the direction of law enforcement. Our findings indicate that this may be an ideal timepoint to assess an individual’s likelihood to progress to more problematic patterns of drug-taking, and to allocate interventional resources. In contrast, assessment of motoric activity at this timepoint might be contaminated by previous drug use, as drugs of abuse (psychostimulants in particular) are known to produce long-lasting plasticity in motor systems (Pierce & Kalivas, 1997). Finally, it is unclear to what extent the HR/HNP models can predict the emergence of negative emotional behavior following drug withdrawal. In fact, in experiments not focused on drug behavior, HR rats are less-prone to depression-like behavior compared to LR controls (Clinton et al., 2011; Stedenfeld et al., 2011). In contrast, we show that animals with high demand for cocaine following the transition to compulsive drug use (but not prior) are more likely to score higher on two depression-like behavioral endophenotypes. Given the strong link between negative emotional state and relapse (Koob & Le Moal, 2008; Koob, 2009), long-term abstinence is likely to be enhanced by identifying those patients most likely to benefit from strategies designed to manage any comorbid depression. In sum, the economic demand approach offers a powerful alternative to the HR/HNP models for predicting addiction vulnerability in individuals with a history of drug use. Future studies that investigate the extent to which the HR, HNP and HD phenotypes overlap with one another, both in terms of behavioral (addiction) and genetic phenotypes, will be of great interest.

### Demand elasticity as a predictor of SB efficacy

We demonstrate that economic demand is also an effective tool for predicting the therapeutic efficacy of the OX1R antagonist SB. We found that demand elasticity for cocaine following LgA, IntA or ShA was directly related to the extent to which SB reduced this measure of motivation for cocaine, such that SB was most effective in high demand animals at both doses tested. Interestingly, animals that fell into the top quartile of demand (lowest α values) were equally responsive to SB10 and SB30, indicating that the orexin system may be particularly strongly engaged in high demand animals. This interpretation aligns with a number of studies that show that SB is more effective at blocking drug seeking in high motivation animals. For example, SB reduces ethanol self-administration and reinstatement selectively in rats that show high preference for alcohol in a two-bottle choice test (Moorman & Aston-Jones, 2009) or greater operant responding for alcohol on an escalating fixed ratio schedule (Moorman et al., 2017). Similar findings have been reported in alcohol-preferring mice using an OX1R antagonist (Lopez et al., 2016), as well as in selectively bred alcohol-preferring rats (Lawrence et al., 2006; Jupp et al., 2011). In addition, we recently reported that SB was only effective at reducing demand for cocaine when motivation was enhanced by the presence of drug-associated conditioned stimuli (Bentzley & Aston-Jones, 2015). These findings align with several studies that have reported that the magnitude of Fos expression in orexin neurons during drug-seeking is positively correlated with drug-seeking behavior (Harris & Aston-Jones, 2006; Hamlin et al., 2007; Richardson & Aston-Jones, 2012; Moorman et al., 2016). Further, we recently showed that animals with high motivation for cocaine following IntA exhibit greater numbers of orexin-immunoreactive cells in lateral hypothalamus (LH) compared to lower-motivation (ShA) controls, and that these LH orexin cells are activated to a greater extent by drug-associated contexts (James et al., in submission). Similarly, pre-pro orexin mRNA is upregulated in LH in alcohol-preferring rats following chronic alcohol consumption (Lawrence et al., 2006). Thus, it is possible that individual differences in the orexin system (cell numbers, receptor densities, receptor binding, cell reactivity and sensitivity to receptor antagonist treatment) might serve as a biomarker of addiction propensity; clearly further work is required to investigate this hypothesis. Regardless, our findings indicate that therapies that block orexin signaling are highly effective in high demand individuals, even at low doses, and that the economic demand task would be a highly effective means to identify individuals most likely to benefit from these therapies. The ability to direct personalized pharmacotherapy to high demand individuals is especially important, as these individuals typically have poor clinical outcomes following psychotherapy-based interventions (Murphy et al., 2015).

### Conclusions

In conclusion, we show that demand elasticity (α), measured using our within-session behavioral economics task after only limited drug-taking experience, is a behavioral biomarker of an individual animal’s propensity to develop several addiction-like endophenotypes following LgA or InTA to cocaine. Re-assessment of demand elasticity after LgA or InTA serves as a strong predictor of future drug-seeking and the emergence of negative emotional behavior during withdrawal. Moreover, the behavioral economic approach can also identify individuals most likely to respond to the anti-addiction properties of an OX1R antagonist. Given that the economic demand model has been validated in cocaine-dependent patients (Bruner & Johnson, 2014), future studies to translate such findings into clinical populations may help guide more effective and efficient treatment for cocaine use disorder.

## Acknowledgements

This work was supported by C.J. Martin Fellowships from the National Health and Medical Research Council of Australia to MHJ (No. 1072706) and HEB (No. 1128089), by a U.S. Public Health Service award from the National Institute of Drug Abuse to GAJ (R01 DA006214), and by the Charlotte and Murray Strongwater Endowment for Neuroscience and Brain Health (GAJ). We would like to gratefully acknowledge Ms. Nikki Koll for her excellent assistance with the behavioral experiments.

## Author Contributions

MHJ, HEB, CMS and GAJ designed the experiments. MHJ, HEB and CMS collected the behavioral data. MHJ analyzed the data. MHJ, HEB and GAJ wrote the paper. All authors provided critical feedback on the manuscript draft.

## Data availability

Data are available upon request from the corresponding author.

## Abbreviations

BE: behavioral economics
SB: SB-334867
LgA: long access
ShA: short access
IntA: intermittent access
HR: high responder
LR: low responder
HNP: high novelty-preferring
LNP: low novelty-preferring
HD: high demand
LD: low demand
ID: intermediate demand

